# Pollinate to Suppress: Sterile Pollen and Certation Effects in Palmer amaranth (*Amaranthus palmeri*)

**DOI:** 10.1101/2025.06.30.662479

**Authors:** Wenzhuo Wu, Mohsen B. Mesgaran

## Abstract

The sterile pollen technique, which involves applying X-ray irradiated pollen to female plants, has shown promising results in reducing seed production in dioecious Palmer amaranth (*Amaranthus palmeri* S. Wats.). However, field-scale implementation of this method requires a carrier for pollen delivery, as applying pure pollen on a large scale is not practical. Additionally, variability in flowering time within and among individual plants may impact the technique’s effectiveness. Mass pollination may also lead to a female-biased sex ratio in the progeny, a phenomenon known as certation. Therefore, in this study, we aimed to: (1) identify a suitable dry (inert) diluent and the most effective pollen-to-diluent ratio for large-scale application, and (2) determine the most effective combination of initiation time, frequency, and number of sterile pollen applications, and 3) test for evidence of certation in *A. palmeri*. Plants were grown in the greenhouse in summer 2021. Sterilized pollen irradiated at 300 Gy was mixed with talc or wheat powder flour, respectively, at six v/v ratios (pollen%/powder%) of 0/100 (powder alone), 5/95, 10/90, 25/75, 50/50, or 100/0 (pure irradiated pollen). An equal amount of the pollen-diluent powder mixture was then brushed onto the standardized lengths of inflorescence of receptive female plants. Flower and seed number on each inflorescence was counted and used to calculate seed set in each treatment. The findings showed that a minimum of 25% irradiated pollen in the mixture, either with talc powder or wheat flour, can effectively reduce seed set in *A. palmeri*.

Building on this, a second round of greenhouse experiments was conducted in fall 2021, where female plants were pollinated with a 25% irradiated pollen and talc powder mixture using a powder duster. Pollination treatments were initiated at different times after anthesis (7, 14, and 21 days), and the number of applications varied (once, twice, or three times) with intervals of one, two, or three weeks between applications. The greatest reduction in seed output was achieved when sterile pollen application began seven days after anthesis and was repeated three times at 7-day intervals. Evidence of certation was also observed in *A. palmeri*, with the progeny of mass-pollinated females showing a higher female ratio (58%) compared to the open-pollinated population (47%). Mass pollination also led to a slight, though not statistically significant, reduction in plant size traits such as height and number of branches, providing an additional potential benefit of the sterile pollen technique.

## Introduction

Weeds can cause considerable damage to agricultural production (Oerke 2006). While herbicides and tillage have been the primary tools used in controlling weeds in modern agricultural systems, over-reliance on these tactics has resulted in negative impacts on both the environment and crop productivity (MacLaren et al. 2020). The breeding system of a weed species plays a critical role in its ability to invade and establish in new environments (Rambuda and Johnson 2004). However, research on reducing seed production in weedy species through exploitation of breeding system limitations has received little attention. Palmer amaranth (*Amaranthus palmeri* S. Wats.), a dioecious weed species with male and female reproductive organs on separate plants, is a highly aggressive and invasive weed species. It is ranked as the worst weed in US corn fields in a survey conducted by the Weed Science Society of America and is now a serious threat to agricultural production systems in over 40 countries (Van Wychen 2020; Webster and Nichols 2012). Dioecy enforces outcrossing, which minimizes inbreeding depression and increases genetic variation within populations through pollen dispersal by male plants (Thomson and Barrett 1981; Charlesworth et al. 1999). Studies have shown that the long-distance dispersal of pollen in *Amaranthus palmeri* has facilitated the transfer of herbicide-resistance genes to susceptible female plants, resulting in offspring with acquired herbicide resistance (Oliveira et al. 2018; Sosnoskie et al. 2012). However, successful fertilization in dioecious species relies on proximity and synchronization of male and female flowers, as well as pollen viability. These limitations present an opportunity for the development of novel management strategies for *A. palmeri*.

One possible strategy for controlling dioecious weed species is to induce pollen sterility, whereby a plant becomes incapable of producing viable seeds. At certain doses, irradiated pollen retains physiological viability and can germinate on the stigma to produce a pollen tube but is genetically inactive and cannot fertilize the egg cell to form seeds (Musial and Przywara 1998). The effectiveness of the sterile pollen technique in managing Palmer amaranth has been shown experimentally, as results showed 300 Gy is the most effective irradiation dose to balance induced sterility and mating competitiveness (Wu and Mesgaran 2021). This dosage significantly reduced seed set by at least 50% when compared to open pollination. This method is similar to the sterile insect technique, which utilizes eco-friendly and species-specific methods to decrease mosquito populations by releasing a large number of sterile male mosquitoes into the environment to mate with females (Parker and Mehta 2007).

This study aimed to explore how to increase the effectiveness of sterile pollen application to reduce seed production in Palmer amaranth. Under field conditions, it can be challenging to uniformly apply small volumes of pollen to stigmas. Dry particulates used as pollen diluents can improve the flow and uniformity of pollen distribution (Desai et al. 1997). Thus, the first objective was to determine an ideal dry (inert) diluent at the most effective mixed ratio for large scale application. Furthermore, due to the indeterminate nature of Palmer amaranth inflorescences (Tranel et al., 2002), flowers are likely to be at various ages when irradiated and sterile pollen is applied (within-individual variation). In addition, not all plants will flower simultaneously, meaning that a proportion of the population will not be exposed to sterile pollen with a single pollination event (between-individual variation). These within-individual and between-individual variations in flowering pattern make it challenging to maximize the efficiency of this technique. Therefore, our second objective was to identify the optimal combination of starting time, frequency, and number of sterile pollen applications to minimize seed production of Palmer amaranth. The aim is to apply the sterile pollen technique with the ideal timing, frequency, and number of applications to result in the minimum possible seed production.

To better understand the ecological implications of sterile pollen technique, we further investigated the effects of mass sterile pollen application. Flower production and seed output are influenced by the development of the inflorescence (Kirchoff et al. 2017). Maximal reproductive success depends on the timing of flowering and on balancing the number of seeds produced with resources allocated to individual seeds (Benlloch et al. 2007). The production of large inflorescences itself can be costly as it may deplete resources that could otherwise be allocated to other plant organs (Suetsugu et al. 2015). A scientific unknown is whether there exists a trade-off between inflorescence growth and fertilization rate. If a trade-off exists, mass pollination with sterile pollen may have the additional benefit of reducing inflorescence growth. Consequently, our next objective was to investigate the effect of mass pollination on inflorescence growth and seed output. We hypothesized that inflorescence growth and total seed output should be reduced by artificial mass pollination (sterile pollen) in Palmer amaranth. Mass pollination may also influence sex ratio due to certation, a prezygotic mechanism of sex determination hypothesized to originate from the competition between a female-determining gamete and a male-determining gamete (Correns 1928). As a result, when a heavy load of pollen is dusted on female flowers, the female-determining gamete would rapidly reach and sire more than half the ovules and leave a smaller proportion of ovules available to male-determining gametes as was found in *Rumex* species (Conn et al. 1981). We hypothesized that mass pollination would change the sex ratio in the progeny population resulting in a female-biased progeny as predicted by certation theory.

## Materials and Methods

### Section 1 - The optimal formulation of the sterile pollen technique

Seeds (10-15) of *Amaranthus palmeri* were planted into 3-L pots filled with UC Davis potting medium containing a ratio of 1 sand:1 redwood sawdust:1 peat in a greenhouse set at 24/32 C night/day temperature regime and extend photoperiod (13-14 hours of lighting). Fertilizers were applied as 80 ml of a general-purpose fertilizer solution (Jack’s Professional General Purpose 20– 20–20, Allentown, PA) weekly at 350 ppm N starting from 2-true leaves with drip irrigation for two minutes and twice per day. Seedlings were thinned to one plant per pot (300 pots in total).

Once plants reached the flowering stage, 100 male and 100 female plants were grown in the same greenhouses to simulate the condition in the field.

Pollen was collected by gently tapping or shaking the male inflorescence, causing the pollen grains to be released onto aluminum foil placed beneath the inflorescence. The collected pollen was then sieved through a 250-mm mesh opening to remove large floral materials and stored in polyvinyl containers at 95 to 100% relative humidity until needed. To sterilize the pollen, fresh and mature pollen was placed in Petri dishes with parafilm and irradiated with gamma-rays from Cesium-137 at the UC Davis Center for Health & the Environment (https://che.ucdavis.edu). The irradiation was delivered at a dosage of 300 Grey (Gy), which was determined to be the most effective irradiation dosage in previous tests (Wu and Mesgaran 2021).

Two types of diluent powders, wheat flour and talc powder, were evaluated for their effectiveness in diluting the sterilized pollen. Previous work has shown that these compounds do not change the biological properties of pollen and can be mixed and applied with pollen uniformly (Wetzstein and Law 1999; Vaknin et al. 1999). The sterilized pollen was mixed with each powder at six v/v ratios (pollen%/powder%) of 0/100 (powder alone), 5/95, 10/90, 25/75, 50/50, or 100/0 (pure irradiated pollen). The pollen-powder mixture was then uniformly applied to the inflorescence, each about 7 inches long (∼18cm), of receptive female plants using the same amount of pollen-diluent mixture. Each treatment had three replications. For each replicate, five 1 cm sections of treated inflorescences were dissected and analyzed to determine the number of flowers, normal full seeds and abnormal seeds (undeveloped ovules or empty seeds). Seed set was calculated by using the number of viable seeds divided by the number of flowers and expressed as percentage. The optimal pollen-powder mixture was determined based on the formulation that resulted in the lowest seed set.

Seed set data from the optimal diluent at the most effective mixed ratio experiment were subjected to dose-response analysis using the DRC package (Ritz et al., 2015) in R software (R Core Team 2019). Because the maximum pollen share in a mixture (%) is 100, a four-parametric log-logistic function (Equation 1; (Streibig et al. 1993) best describes seed set in relation to the pollen share in mixture (%) of talc and wheat powder :

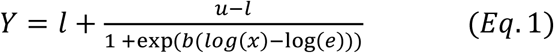

where *Y* is seed set, *e* is the effective pollen share in a mixture producing a seed set halfway between the *u* and 𝑙 parameters, 𝑢 is the upper limit, 𝑙 is the lower limit of the curve, and *b* is the relative slope around the inflection point (*e*). Data from the two different powders were pooled because there was no difference between the full model and the reduced (pooled) model.

### Section 2 - The optimal application of the sterile pollen technique

The optimal sterile pollen-powder mixture identified in section one was used for this subsequent experiment. Plant material and pollen collection remained consistent with the methods previously described. Based on preliminary results, it was estimated that 1 ml of pollen contains approximately 5,000,000 individual grains. Furthermore, our observations indicate that one centimeter of inflorescence contains approximately 100 flowers on average. Assuming that approximately 20% of the pollen is expected to be distributed onto female plants, the volume of pollen needed for each experiment was calculated as follows:

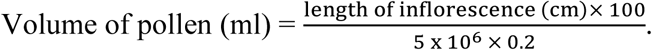

Female plants were pollinated with sterile pollen and powder mixture at different starting times after anthesis (when we can visually identify the sex of plants), which includes 7 days, 14 days, and 21 days. The number of applications were once, twice and three times. Application time intervals were one week, two weeks and three weeks. Open pollination was used as control. Each combination of treatments (see Figure 1) had three replicates. Pollination was done with a powder duster. The duster was squeezed to release the powder and then the nozzle applicator guided the powder to targeted female flowers. After seeds matured, plants were crushed and sieved to collect seeds. The total amount of seeds was weighed; five groups of 100 seeds were weighed and averaged to calculate the total number of seeds produced.

**Figure 1.**
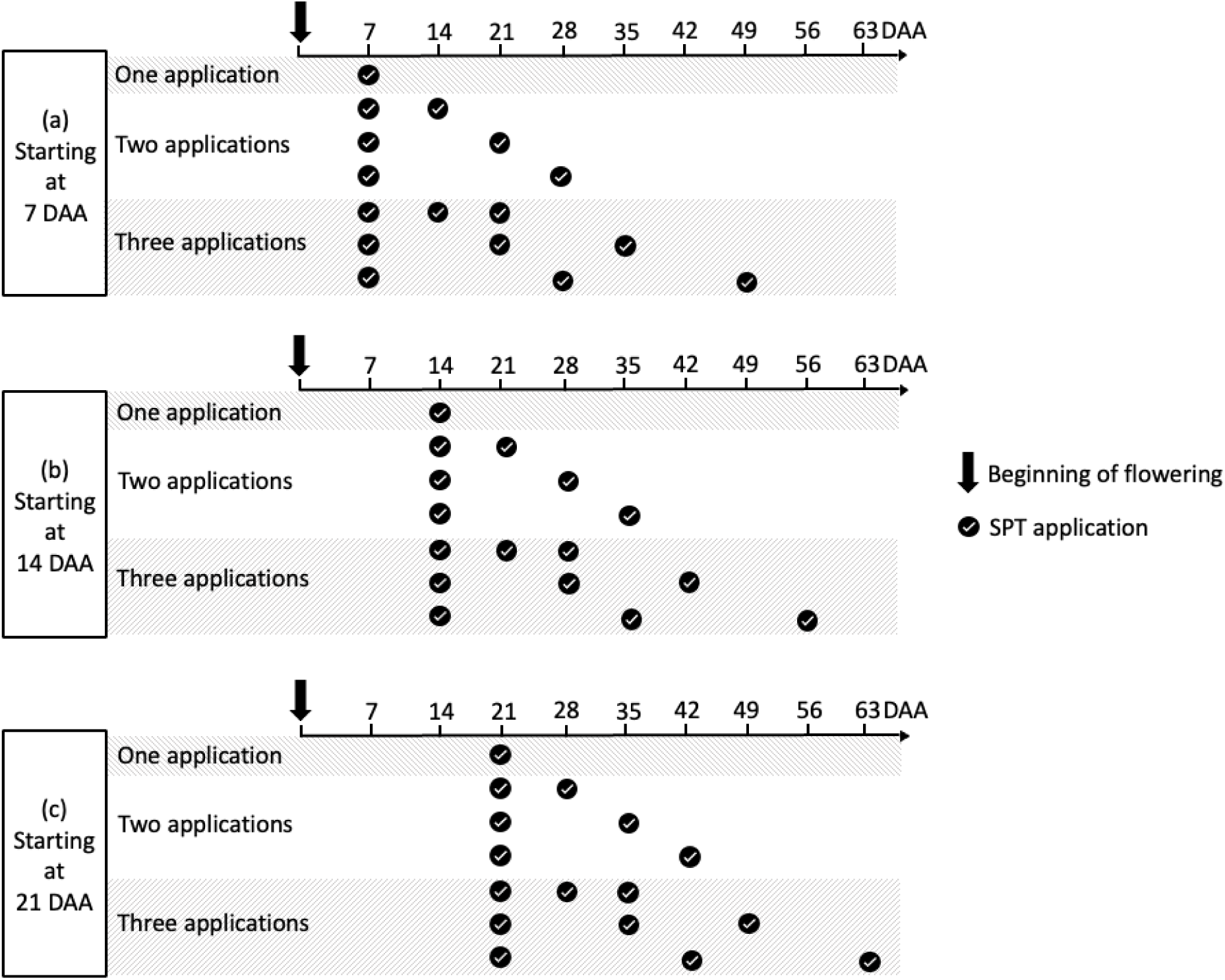
Application plan of sterile pollen technique (SPT). DAA = days after anthesis. (a) When the application starts at 7 DAA, it was applied once, twice at 7-day, 14-day, and 21- day intervals, and three times with 7-day, 14-day, and 21-day intervals. (b) When the application starts at 14 DAA, it was applied once, twice at 7-day, 14-day, and 21- day intervals, and three times with 7-day, 14-day, and 21-day intervals. (c) When the application starts at 21 DAA, it was applied once, twice at 7-day, 14-day, and 21- day intervals, and three times with 7-day, 14-day, and 21-day intervals.

Since the optimal application of the sterile pollen technique experiment is not a full-factorial design, three factors: application starting time, the number of applications and application intervals were combined into a single factor and a one-way ANOVA was performed by using aov() functions in R (R Core Team 2019). Number of seeds estimated per plant from the treatments was compared to the seed production from open pollination using ANOVA with Dunnett’s test.

### Section 3 - The effect of mass pollination on inflorescence growth and certation

In this experiment, we employed the optimal application scheme identified in section two but with mass pollen. Plant material and pollen collection are the same as described previously. To achieve a mass pollination effect, we utilized a pollen quantity that was one hundred times greater than the standard amount:

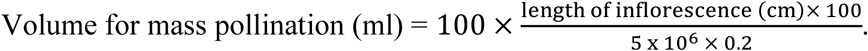

After anthesis, male and female plants were randomly separated into three different greenhouses for specific treatments: open pollination (control), pure mass irradiated pollen and pure mass non-irradiated pollen. Each treatment was replicated 5 times and applied using brushes. Greenhouse-one served as the control with five females and five males (open pollination). Greenhouse-two had five females and five males with females receiving mass sterile pollen based on the optimal application scheme identified in section two. Similarly, greenhouse-three had five females and five males, with females receiving mass fertile pollen following the same optimal application scheme. These two greenhouses had both female and male plants to simulate the condition in the field. An additional 50 male plants were kept outdoors for pollen collection.

Once seeds reached maturity, we assessed various aspects of inflorescence outgrowth, including plant height, the number of branches per plant, length of the main inflorescence, total length of all inflorescences, and dry weight. Following these measurements, the plants were crushed and sieved to collect the seeds. The total seed weight was recorded, and we calculated the total seed production by weighing and averaging five groups of 100 seeds. A t-test was used to compare the differences in inflorescence growth and seed production for treatments using mass, irradiated pollen or non-irradiated pollen against the results from open pollination. This analysis was conducted using the R software.

Lastly, seeds from the mass pollination experiment were used for a certation experiment.

Two hundred seeds were randomly selected from female plants within the same treatment and were then planted in the greenhouse under the previously described conditions. Plant sex was recorded after anthesis. Finally, the sex ratio observed in both the irradiated mass pollination group and the non-irradiated mass pollination group was subjected to a statistical analysis for comparison against the sex ratio found in the open pollination group using a chi-square test.

## Results and Discussion

### Section 1 - Optimal formulation of the sterile pollen technique

Two powder types were effective in reducing the seed set in *Amaranthus palmeri* as shown by our dose-response analysis (Figure 2). Data from the two tested powder types were pooled because there was no difference between the full model (with the powder type as a covariate) and the reduced model (without the powder type as a covariate), as shown in Appendix 1. This suggests four parameters (𝑒, 𝑙, 𝑢, 𝑏) can be fixed across curves of talc powder and wheat powder (flour) without significantly reducing the goodness of fit. Under the competition with naturally-occurring pollen, seed set was 50.73% and 28.03% when pure powder and pure irradiated pollen were applied, respectively, as indicated by the values of parameters 𝑢 and 𝑙 (Table 1). The seed set in both treatments was lower than the seed set from open pollination (64.15% ± 4.52%). The effective pollen share in the mixture was 4.81 (Table 1), producing a seed set halfway between the lower limit and upper limit. The ED_50_, the ratio reducing seed set by 50%, was not estimable because the lower limit of the model was greater than half the maximum response (i.e. 50.73% × 0.5 = 25.365%) (Keshtkar et al. 2021). To minimize seed set and conserve irradiated pollen, a mixture ratio of 25%:75% should be used; this is the smallest ratio that yielded seed set close to the lower limit (28.03%) while minimizing the amount of irradiated pollen required (Figure 1). Therefore, using a 25%:75% mixture ratio can effectively reduce seed set in *A. palmeri* while conserving the limited resources of irradiated pollen.

**Figure 2.**
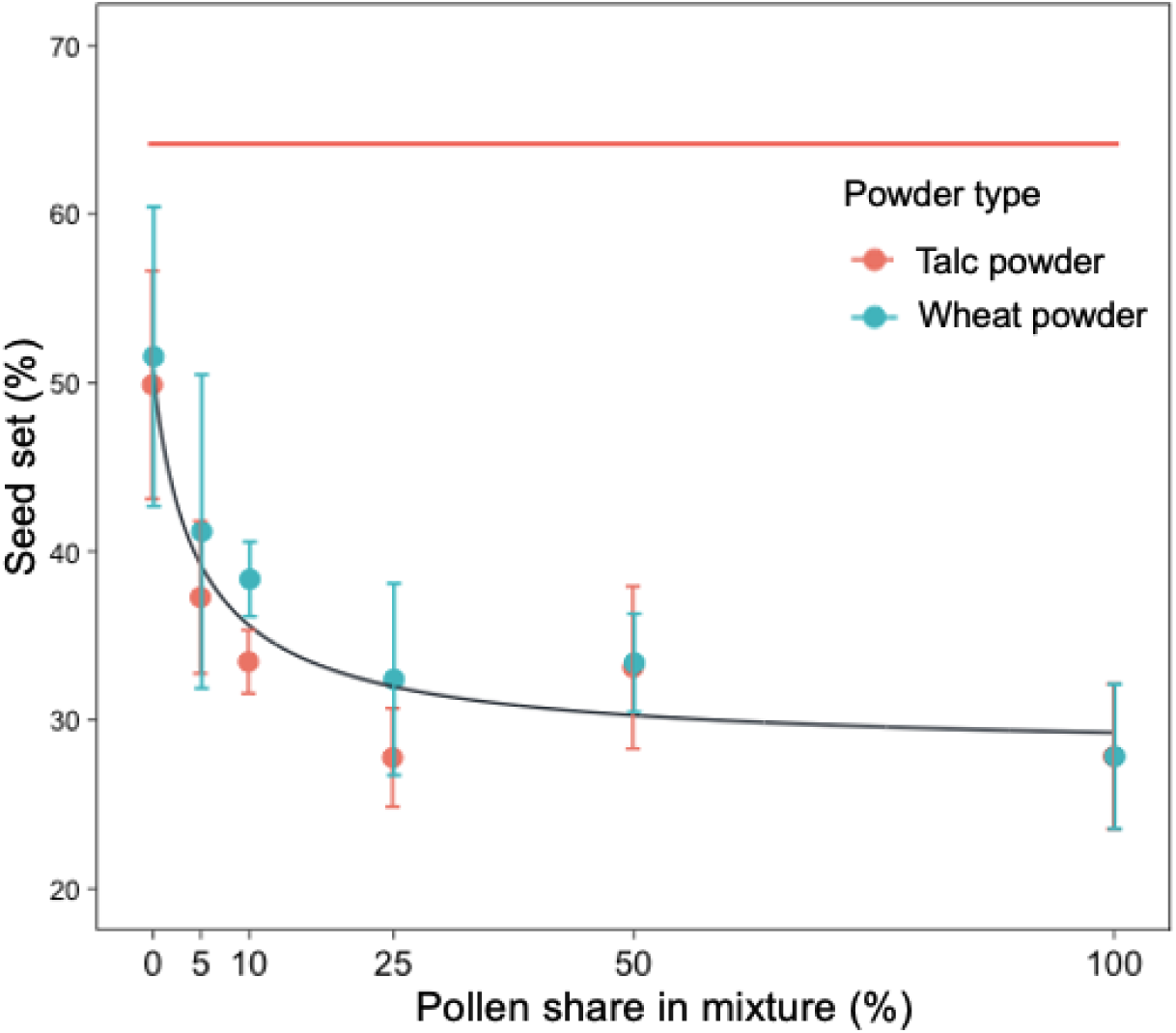
Dose response curve showing seed set in Palmer amaranth at different pollen shares in mixture (%) with talc powder and wheat powder flour (Equation 1) under greenhouse conditions. Black line is fitted value of pooled data, and red flat line represents the average seed set for open pollination. Solid circles indicate observed seed set averaged with three replications each. Error bars indicate standard error. Model parameter estimates are shown in Table 1. Note: To minimize seed set and conserve irradiated pollen, a pollen share in mixture of 25% should be used; this is the smallest ratio that yields a seed set close to the lower limit (28.03%) while minimizing the amount of irradiated pollen required.

**Table 1.** Estimated parameter values for the four-parameter (Equation 1) log-logistic models used to describe seed set in Palmer amaranth in response to increasing irradiated pollen share in mixture with talc powder and wheat powder in greenhouse conditions.

| Parameter | Estimate | Std. Error | t-value | p-value<sup>b</sup> |
| --- | --- | --- | --- | --- |
| $b^a$ | 0.9434 | 1.5329 | 0.6155 | 0.5626 |
| $l^a$ | 28.0285 | 8.9542 | 3.1302 | 0.0037** |
| $u^a$ | 50.7323 | 3.4407 | 14.7448 | <0.0001*** |
| $e^a$ | 4.8075 | 3.8732 | 1.2412 | 0.2235 |

Residual standard error: 0.0844 (32 degrees of freedom).
<sup>a</sup>The parameters are fixed across different diluents. $e$ is the effective pollen share in a mixture producing a seed set halfway between the $u$ and $l$ parameters, where $u$ is the upper limit, $l$ is the lower limit of the curve and $b$ is the relative slope around the inflection point ( $e$ ).
<sup>b</sup>Statistical significance is denoted by asterisks (\* for $p < 0.05$ , \*\* for $p < 0.01$ , \*\*\* for $p < 0.001$ ).

The lower seed set observed with pure powder application compared to open pollination can be attributed to the physical barrier created by the powder, which covers the stigma and prevents non-irradiated pollen from fertilizing the ovule and producing seeds (Vaknin et al 1999). With an increase in the proportion of irradiated pollen in the mixture, there is a decrease in the seed set. This is due to the fact that while the irradiated pollen can germinate on the stigma and produce a pollen tube, it is incapable of fertilizing the egg cell, thereby failing to produce any seeds (Musial and Przywara 1998).

When selecting an optimal diluent for pollen application, it is crucial to take into account factors such as non-toxicity and preventing any disruption to pollen-stigma interactions. Artificial supplementary application of pollen using non-toxic diluents like wheat flour and talc powder, has shown positive results in various plants. For instance, in raspberry (*Rubus idaeus* L.), talc-diluted pollen is employed to enhance fruit production (Jennings and Topham 1971). Similarly, in *Cannabis sativa*, the use of cryopreserved pollen mixed with wheat flour yields seeds of comparable number, size, and morphology to those produced with untreated fresh pollen (Gaudet et al 2020). However, when wheat flour is combined with irradiated pollen of Palmer amaranth, it tends to clump and degrade pollen flow more than talc powder. This is due to wheat flour’s higher water absorption capacity 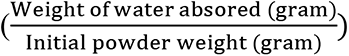 which is approximately 80%, compared to the 1.5% for talc powder (Vaknin et al. 1999; Yi et al. 2003). Additionally, the size of the diluent particles should be similar to that of the pollen to achieve consistent mixing and uniform application. Talc powder has a median diameter of 26.57 μm whereas milling produces wheat flour particles with a median diameter of about 120 µm (Gilbert et al 2018; Liu et al 2016). The diameter of Palmer amaranth pollen is approximately 31 µm (Wu et al. 2023). Therefore, talc powder is a more suitable diluent than wheat flour.

### Section 2 - Optimal application of the sterile pollen technique

The combined effect of sterile pollen application starting time, application frequency and application interval had a statistically significant impact on seed production (Figure 3, Appendix 2). Initiating application 7 days after anthesis consistently resulted in the lowest estimated seed production per plant, significantly outperforming open pollination (P<0.001) as shown in Figure 3. Estimated seed production from single applications at 14 or 21 days after anthesis did not show a significant difference compared to open pollination (Figure 3). Regardless of the start time, increasing the number of applications or reducing the interval between them further decreased seed production per plant. Based on these findings, the optimal application strategy for the sterile pollen technique is to begin at 7 days after anthesis and apply three times at 7-day intervals.

**Figure 3.**
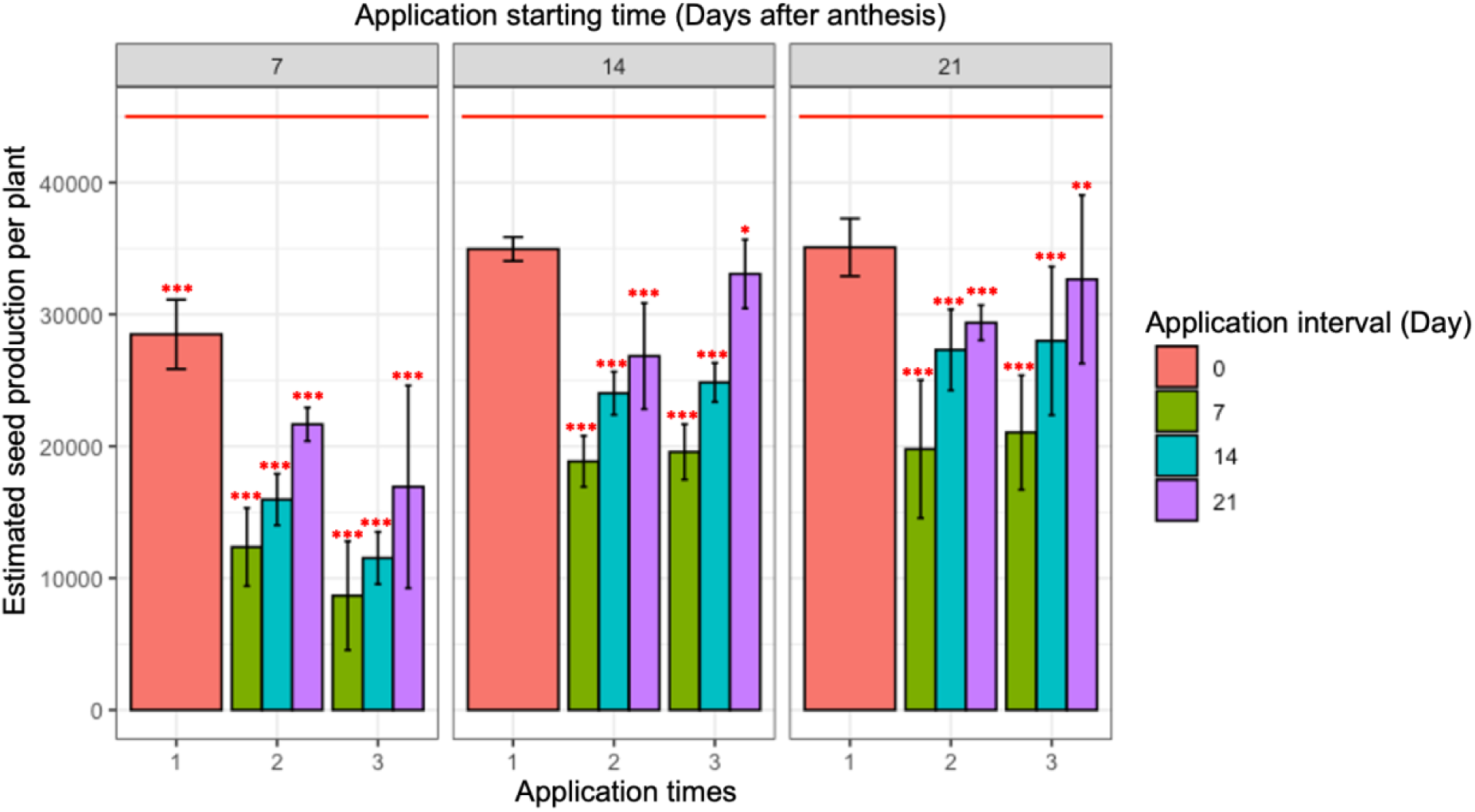
Average seed production per plant in Palmer amaranth, measured across three replications for each combination of application starting time, interval, and times, in greenhouse condition. Error bars indicate standard errors. The red solid line represents the average seed production per plant for open pollination. Significance levels are denoted by asterisks (* for p < 0.05, ** for p < 0.01, *** for p < 0.001) based on Dunnett’s test for comparing treatments with open pollination (control). Note: Seed count per plant was significantly reduced in sterile pollen application compared to open pollination, with the exception of single applications made at either 14 or 21 days after anthesis.

Due to the indeterminate nature of Palmer amaranth inflorescences (Tranel et al. 2002), flowers varied in age at the time we applied irradiated and sterile pollen, resulting in within-individual variation. Additionally, not all plants within the population flower simultaneously, leading to between-individual variation where a portion of the population may not be exposed to sterile pollen. Those flowering variations among female flowers present challenges in terms of the timing for the application of the sterile pollen technique. Furthermore, the initiation of flowering occurred earlier in males than females under both water stress and control conditions (Mesgaran et al. 2021). When applying a pollen-powder mixture to female flowers, our aim is to cover as many stigmas as possible while minimizing the influence of naturally occurring pollen on seed development, considering the earlier flowering of males compared to females. Through our experiments, the optimal application strategy we identified above allows us to achieve minimal seed production while using a reduced amount of irradiated pollen.

The time of flower opening in Palmer amaranth marks the onset of a period in which pollen will be released from male flowers and when pollination, fertilization, and seed production occur in female flowers. Therefore, the timing of flower opening in females plays a significant role in determining the appropriate initiation and interval for application of the sterile pollen technique. In Palmer amaranth, the flowers are open continuously. Flower opening behavior in Palmer amaranth can be influenced by factors such as the time of day and the position of the flower within the inflorescence (van Doorn and Van Meeteren 2003). Based on our observations, it appears that flowers situated in the middle lower part of the inflorescence tend to open first. In many species, including Palmer amaranth, flower opening occurs in the morning, correlated with an increase in temperature and light intensity, and with a decrease in ambient humidity (Mondo et al 2022).

The majority (62 to 73%) of plant species, as indicated by studies utilizing the GloPL Dataset (Lebot et al. 2019), the Konstanz Breeding System Dataset (Agre et al. 2020), and the Stellenbosch Breeding System Dataset (Mondo et al. 2021), demonstrate pollination frequency plays a critical role in determining seed production (Ashman et al 2004; Bennett et al 2020). The reduced seed-set rate in early-flowering plants was associated with pollen limitation, as observed in species like *Peucedanum multivittatum* and *Rhododendron aureum* (Kudo, 1993; Kudo and Hirao, 2006). While the failure of pollen tubes to enter ovules is a common cause of reduced seed production, it is not the sole factor contributing to low fertility, as post-fertilization ovule abortion has been observed to decrease fertility in alfalfa (Sayers and Murphy, 1966).

Additionally, reports indicate that embryo abortion can also lead to reduced seed production in various plant species, including red clover (*Trifolium pratense*) and garden pea (*Pisum sativum*), as well as in plant families other than the Fabaceae (Sato 1956; Linck 1961; Brink and Cooper 1947).

### Section 3 - The effect of mass pollination on inflorescence outgrowth and certation

Regarding mass pollination with non-irradiated and fertile pollen in Palmer amaranth, although there were no statistically significant differences in plant height, branch number, inflorescence length, dry weight, seed weight, and seed production per plant compared to open pollination (Appendix 3), there was a trend suggesting that mass pollination might lead to reductions in these attributes, as shown in Figure 4. This indicates our assumed tradeoff between inflorescence outgrowth and fertilization rate is not significant. Regarding mass pollination with irradiated and sterile pollen in Palmer amaranth, there was a significant reduction in both seed weight and seed production per plant compared to open pollination, as shown in Figure 4. This reduction is likely due to the sterility of the pollen. Other plant attributes did not show significant differences when compared to open pollination. These results suggest that when plants receive a substantial amount of sterile pollen, the presumed trade-off between inflorescence growth and fertilization rate is not significant either.

**Figure 4.**
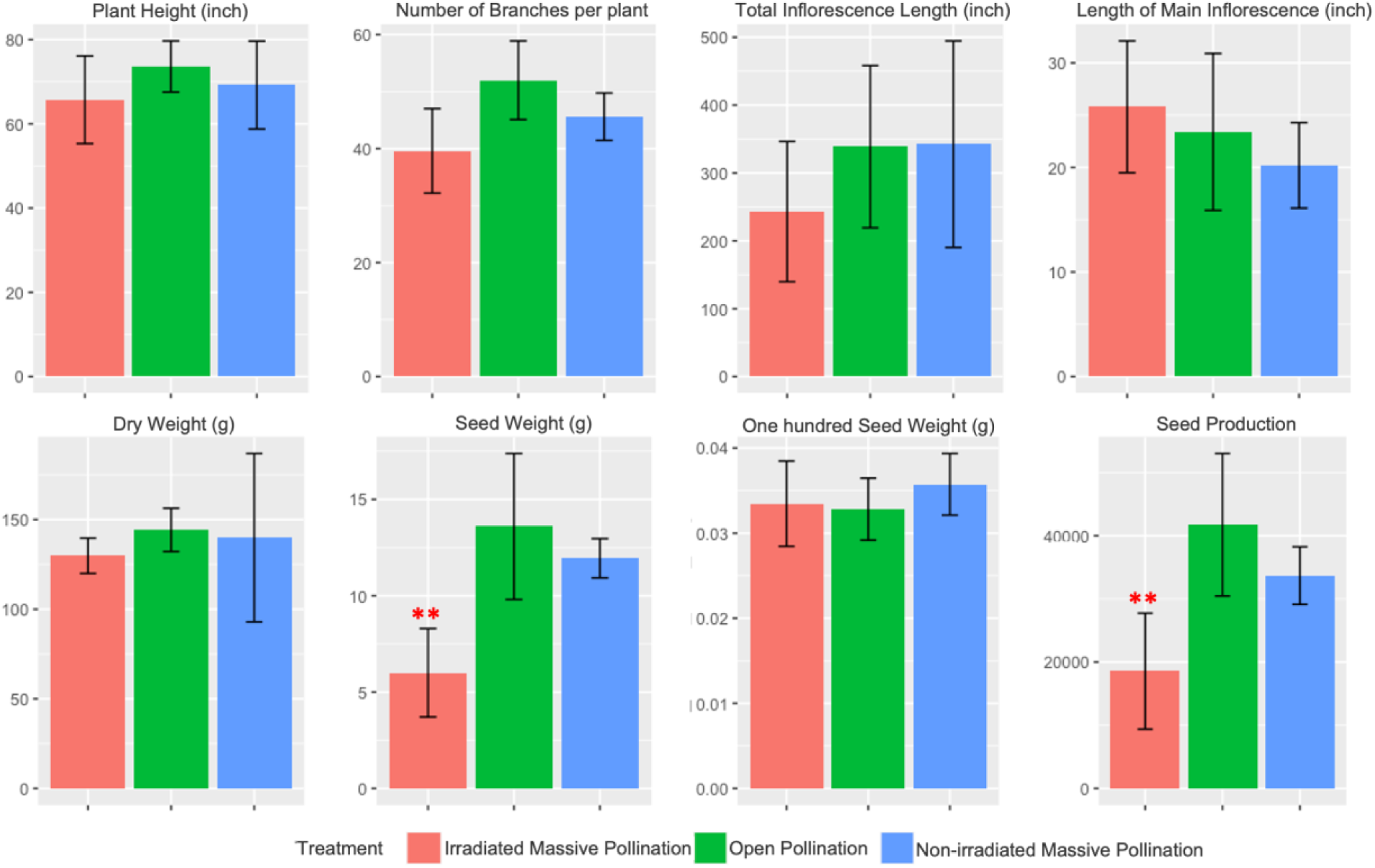
Average values for inflorescence growth measurements and seed production per plant of palmer amaranth, with five replications. Error bars represent standard errors. Significance levels are indicated by asterisks (* for p < 0.05, ** for p < 0.01, *** for p < 0.001), based on t- tests comparing each treatment with open pollination (control). Note: Seed weight and seed count significantly decreased following mass pollination with irradiated and sterile pollen, in comparison to open pollination.

The activity and development of apical and lateral buds, as well as fruits, are controlled by light, temperature, hormone, sugar, and nutrient signaling (Montgomery, 2008; Barbier et al., 2019). These signals enable communication between the shoot apex and lateral sinks (meristems or fruits), ensuring the plant’s architecture and reproductive capacity align with available resources (Walker and Bennett, 2018; Barbier et al. 2019). In annual plants, the suppression of inflorescence growth due to fruit load typically occurs at the late stage of inflorescence development, referred to as the end of the flowering transition (Gonzalez-Suarez et al. 2020; Ware et al. 2020). For instance, in Arabidopsis (*Arabidopsis thaliana*), during this phase, the inhibition of inflorescence shoot growth by fruit load is regulated by auxin and carbohydrate signaling (Goetz et al. 2021). Our findings suggest that the rate of fertilization has a minor effect on inflorescence outgrowth in Palmer amaranth, likely because the development of fruit in Palmer amaranth, which is a thin membranous structure known as an utricle, has relatively low costs.

Pollen limitation has two aspects: quality limitation and quantity limitation. Quality limitation refers to the reduced effectiveness of pollination due to the inferior quality of pollen. In Palmer amaranth, irradiated pollen under 300 Gy doses is genetically inactive and cannot fertilize the egg cell to form seeds (Wu and Mesgaran 2021). In addition, much literature on inbreeding depression has shown that pollen quality effects associated with both self-fertilization and mating between related plants (Herlihy and Eckert 2004) can also reduce seed production (Charlesworth and Charlesworth 1987). This reduction is likely because embryos homozygous for deleterious alleles die during development. On the other hand, traditional pollen limitation is typically associated with plants receiving an insufficient quantity of pollen grains to fertilize all their ovules (quantity limitation). Extensive reviews show that supplemental pollination often increases (Burd 1994, Ashman et al. 2004), and rarely decreases (Young and Young 1992), seed or fruit production.

Regarding sex ratio in offspring after mass pollination in Palmer amaranth, results from mass pollination with irradiated and non-irradiated consistently showed the sex ratio in the progeny population is female dominant as predicted by certation theory (Table 2). It is a prezygotic mechanism of sex determination hypothesized to originate from the competition between a female-determining gamete and a male-determining gamete (Correns, 1928). As a result, when a heavy load of pollen is dusted on female flowers, the female-determining gamete would rapidly reach and sire more than half the ovules and leave a small proportion of ovules available to male-determining gametes as was found in *Silene alba* (Taylor, 1996) and *Rumex* species (Conn et al. 1981). Several other mechanisms have been proposed to account for female bias. In species with sex chromosomes where males are heterogametic, Y-chromosome degeneration may lead to female-biased populations (Smith 1963). This is due to sex viability differences and sex-chromosomal genotype performance during pollination and fertilization. Studies on *Rumex nivalis*, a species with heteromorphic sex chromosomes (XX in females, XY_1_Y_2_ in males), show that both certation and gender-based mortality contribute to female-biased sex ratios (Stehlik et al. 2007, 2008). Research using sex-specific markers across different life stages revealed that female bias starts in pollen and intensifies from seeds to flowering.

**Table 2.** Observed counts of female and male Palmer amaranth plants in the progeny resulting from open pollination, non-irradiated mass pollination, and irradiated mass pollination. We conducted chi-square tests to compare each of the two treatments with open pollination separately.

|  | Female | Male | Treatment<br>comparison | Chi-Square<br>value | P-value<sup>a</sup> |
| --- | --- | --- | --- | --- | --- |
| Open pollination | 94 | 106 |  |  |  |
| Non-irradiated mass pollination | 117 | 83 | Open vs non-irradiated | 4.8547 | 0.0276* |
| Irradiated mass pollination | 113 | 87 | Open vs irradiated | 3.244 | 0.0717 |

<sup>a</sup>Statistical significance is denoted by asterisks (\* for p < 0.05, \*\* for p < 0.01, \*\*\* for p < 0.001).

Environmental factors, like proximity of females to males, affect these ratios (Stehlik et al. 2008); females nearer to males capture more pollen, resulting in more female-biased ratios. Experiments confirm that higher pollen loads intensify this bias (Stehlik and Barrett 2006), supporting Correns’ certation hypothesis that larger pollen loads increase gametophytic competition, favoring fertilization by female-determining pollen tubes.

In conclusion, we investigated how to improve the efficiency of the sterile pollen technique for reducing seed production in *A. palmeri* under greenhouse conditions. The results demonstrated that talc powder is a more suitable diluent than wheat flour. The optimal formulation is to utilize a mixture of 25% irradiated pollen to 75% talc powder by volume, which enhances pollen distribution by improving flow and uniformity. Furthermore, the efficiency of the sterile pollen technique is affected by variations in the timing of female flower opening and interference from naturally-occurring pollen. These factors make it challenging to further enhance the technique’s efficiency. However, through our investigations, we identified the optimal sterile pollen application strategy: initiating application 7 days after anthesis and repeating it three times at 7- day intervals. This strategy allows us to achieve reduction in seed production while also minimizing the amount of irradiated pollen required. Lastly, we found mass pollination of irradiated pollen or non-irradiated pollen did not have an effect on inflorescence growth, but it did affect sex ratio in the progeny population, resulting in slightly female-biased progeny as predicted by certation theory.

## Appendix

**Appendix 1.** Comparison of full model (with powder type as a covariate) with reduced model (without powder type as a covariate) using anova() functions in R for dose response of seed set in Palmer amaranth at different pollen share in mixture (%) with talc powder and wheat powder under greenhouse conditions.

| Model | Model-DF | RSS | DF | F-value | p-value |
| --- | --- | --- | --- | --- | --- |
| Full model | 32 | 2277.6 |  |  |  |
| Reduced model | 28 | 2183 | 4 | 0.3032 | 0.8733 |

**Appendix 2.**
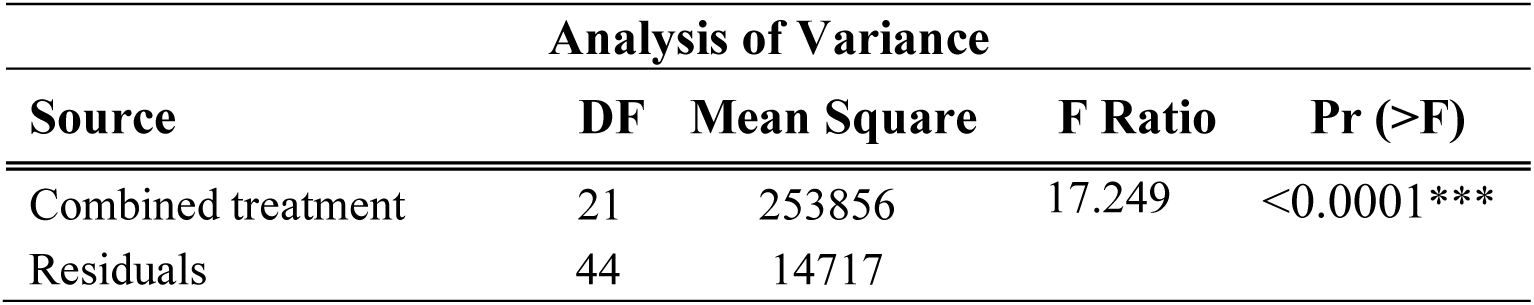
ANOVA table of the effect of combined factors of application starting time, interval, and times on seed production per plant in Palmer amaranth. The three factors, application starting time, the number of applications and application intervals were combined into a single factor and a one-way ANOVA was performed by using aov() functions in R.

**Appendix 3.** T-tests were performed to compare inflorescence outgrowth measurements and seed set per plant of Palmer amaranth, with five replications, between each treatment (irradiated mass pollination and non-irradiated mass pollination) and open pollination (control).

| Comparison | Plant Features | T-Statistic | P-Value<sup>a</sup> |
| --- | --- | --- | --- |
| Irradiated mass pollination<br>vs.<br>Open pollination | Plant height | -1.46 | 0.189 |
|  | Total length of inflorescence (inch) | -0.26 | 0.801 |
|  | Length of main inflorescence (inch) | 1.08 | 0.313 |
|  | Number of branches | -1.06 | 0.801 |
|  | Dry weight (g) | -2.07 | 0.073 |
|  | Seed weight (g) | -3.83 | 0.007** |
|  | One hundred seed weight (g) | 0.23 | 0.823 |
|  | Seed production | -3.56 | 0.007** |
| Non-irradiated mass pollination<br>vs.<br>Open pollination | Plant height | -0.81 | 0.443 |
|  | Total length of inflorescence (inch) | 1.07 | 0.325 |
|  | Length of main inflorescence (inch) | -0.31 | 0.762 |
|  | Number of branches | -1.77 | 0.121 |
|  | Dry weight (g) | -0.20 | 0.850 |
|  | Seed weight (g) | -0.94 | 0.393 |
|  | One hundred seed weight (g) | 1.27 | 0.239 |
|  | Seed production | -1.47 | 0.196 |

<sup>a</sup>Significance levels are indicated by asterisks (\* for $p < 0.05$ , \*\* for $p < 0.01$ , \*\*\* for $p < 0.001$ ). The decrease in seed weight and seed production were statistically significant in mass pollination with irradiated and sterile pollen, compared to open pollination.

## Notes

### Competing Interest Statement

The authors have declared no competing interest.

### Funding

- USDA-NIFA: CA-D-PLS-2511-H

